## Supplementary figures and images for "Omics of an enigmatic marine amoeba uncovers unprecedented giant viruses gene trafficking and provides insights into its complex life cycle"

### Supplementary Figures

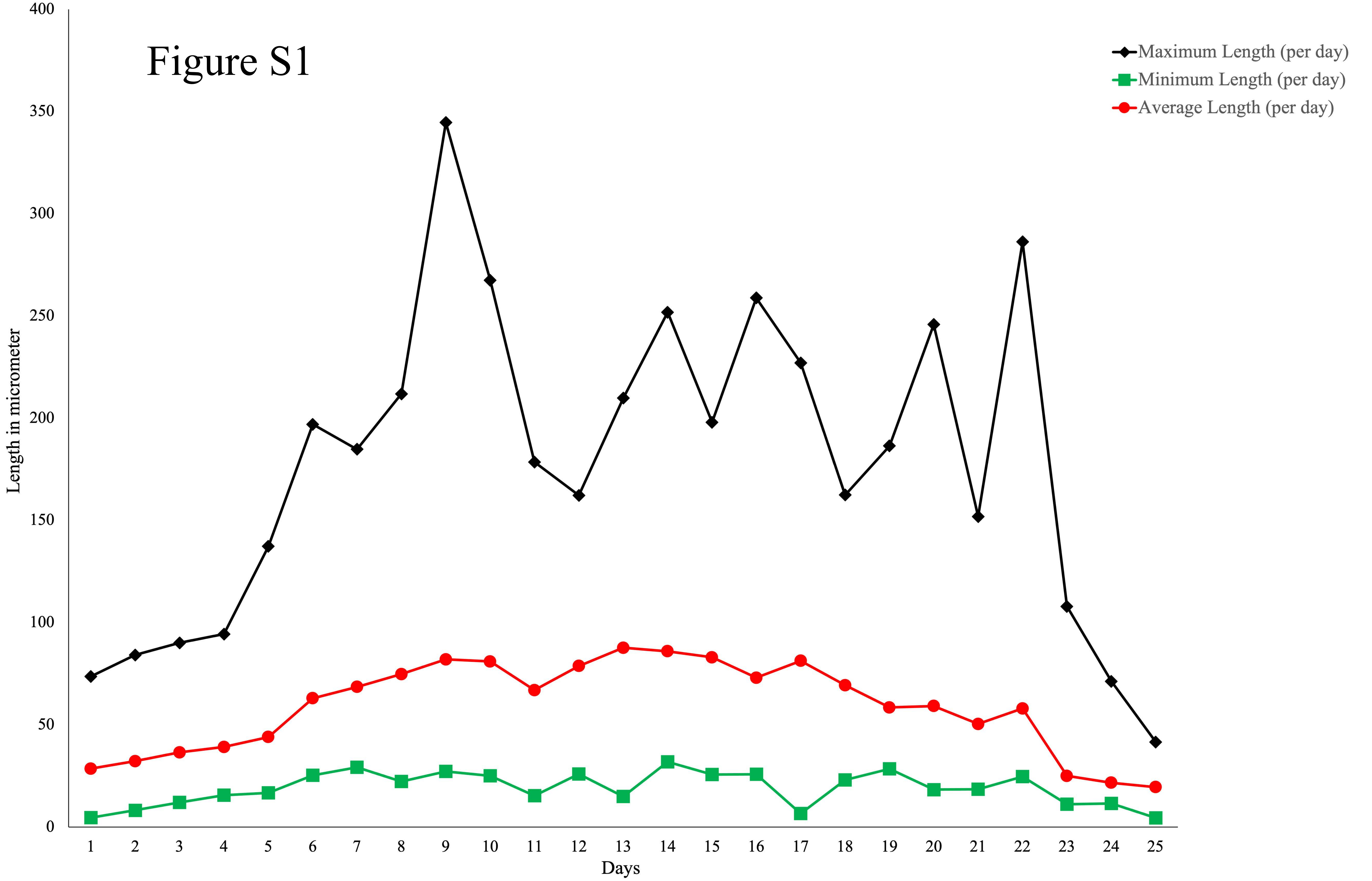

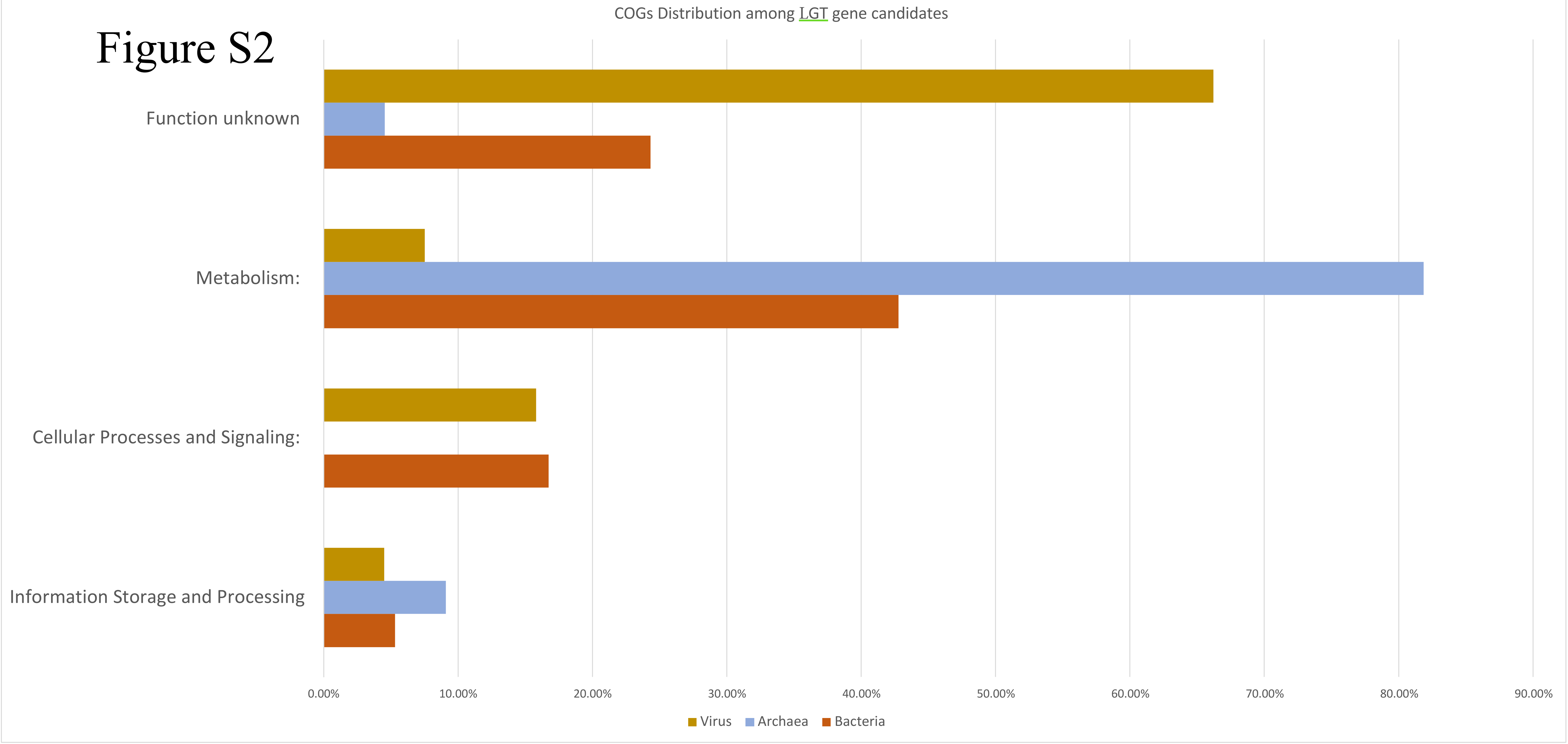

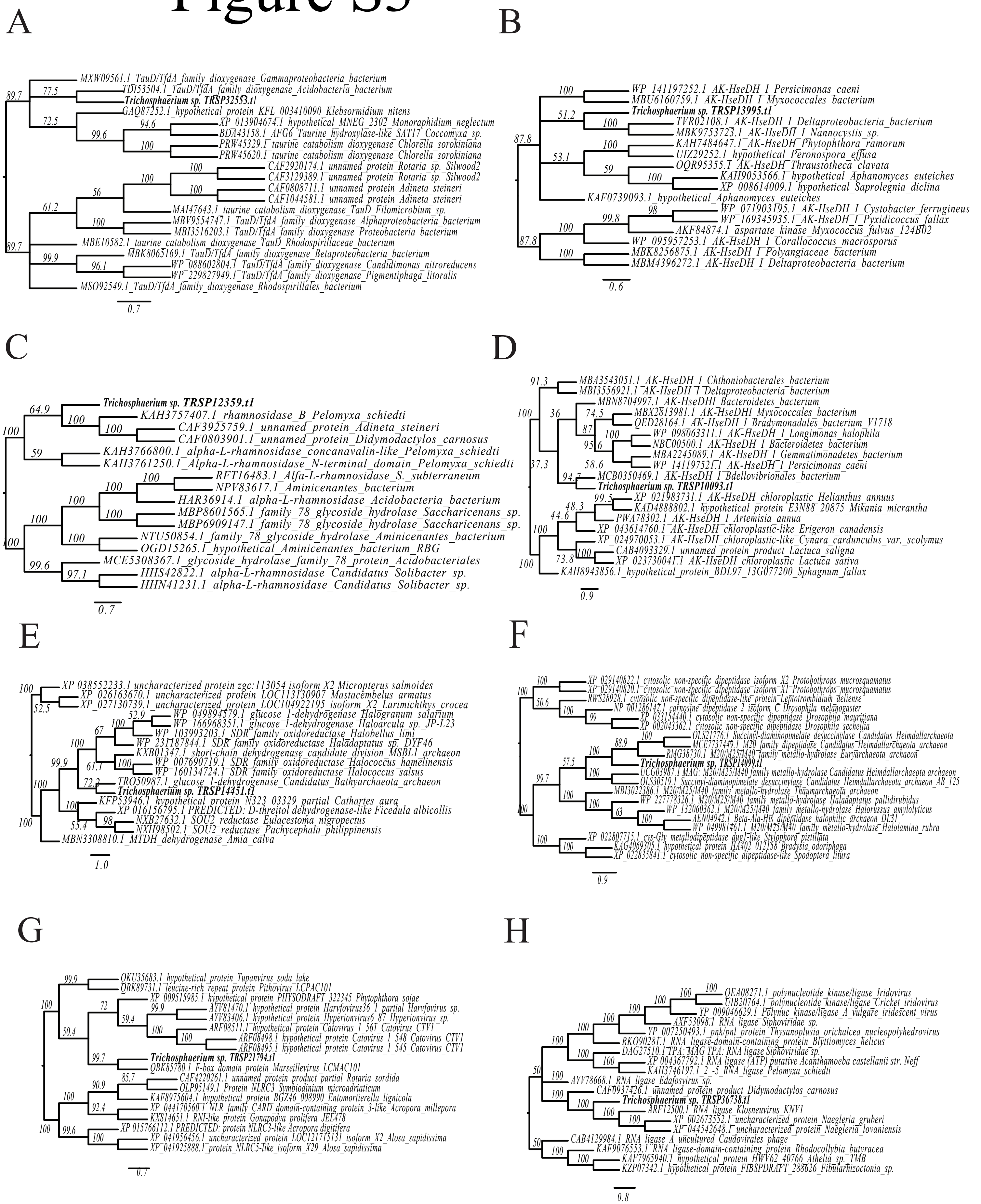

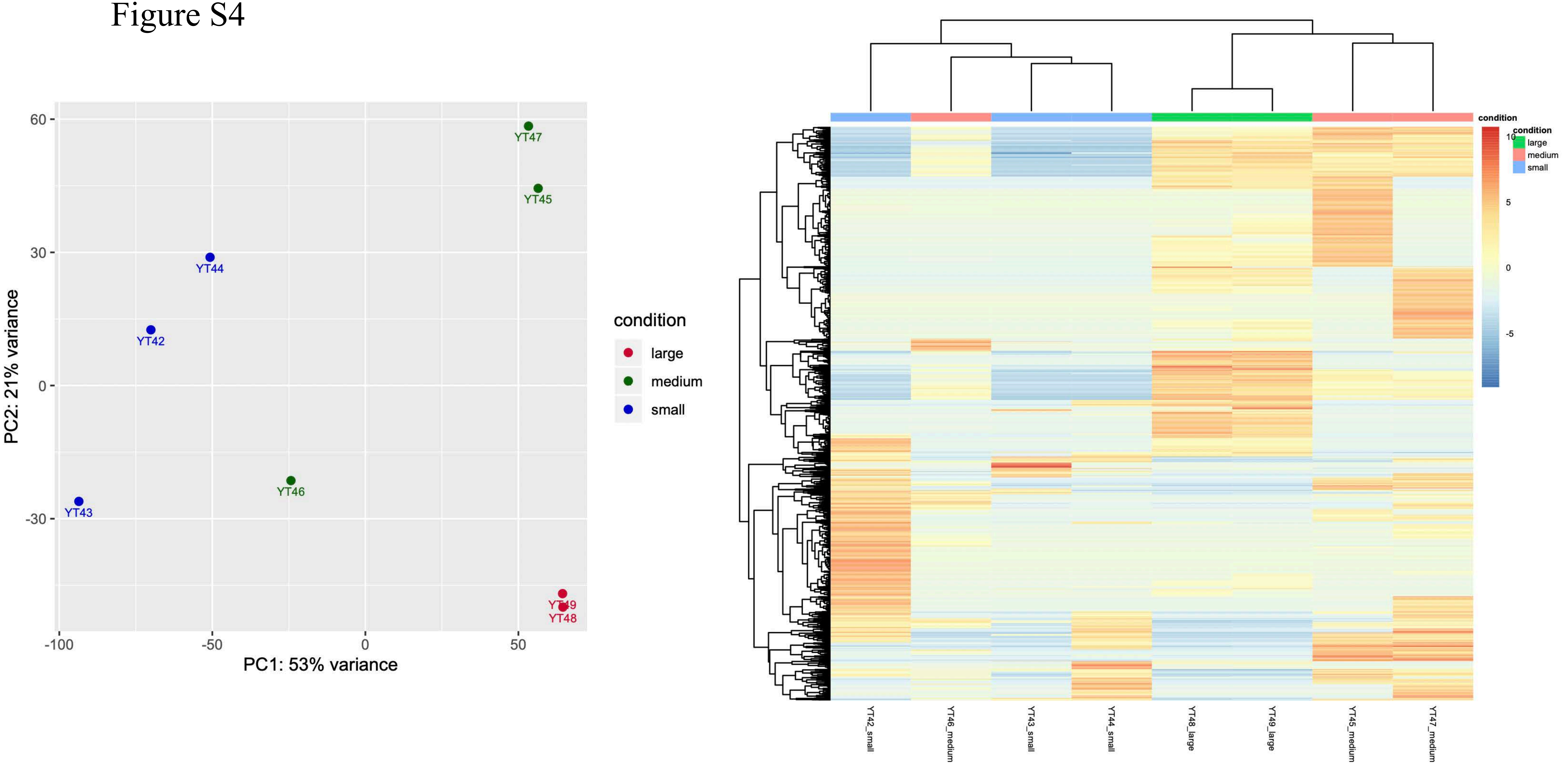

Figure S5

#### PCA plot

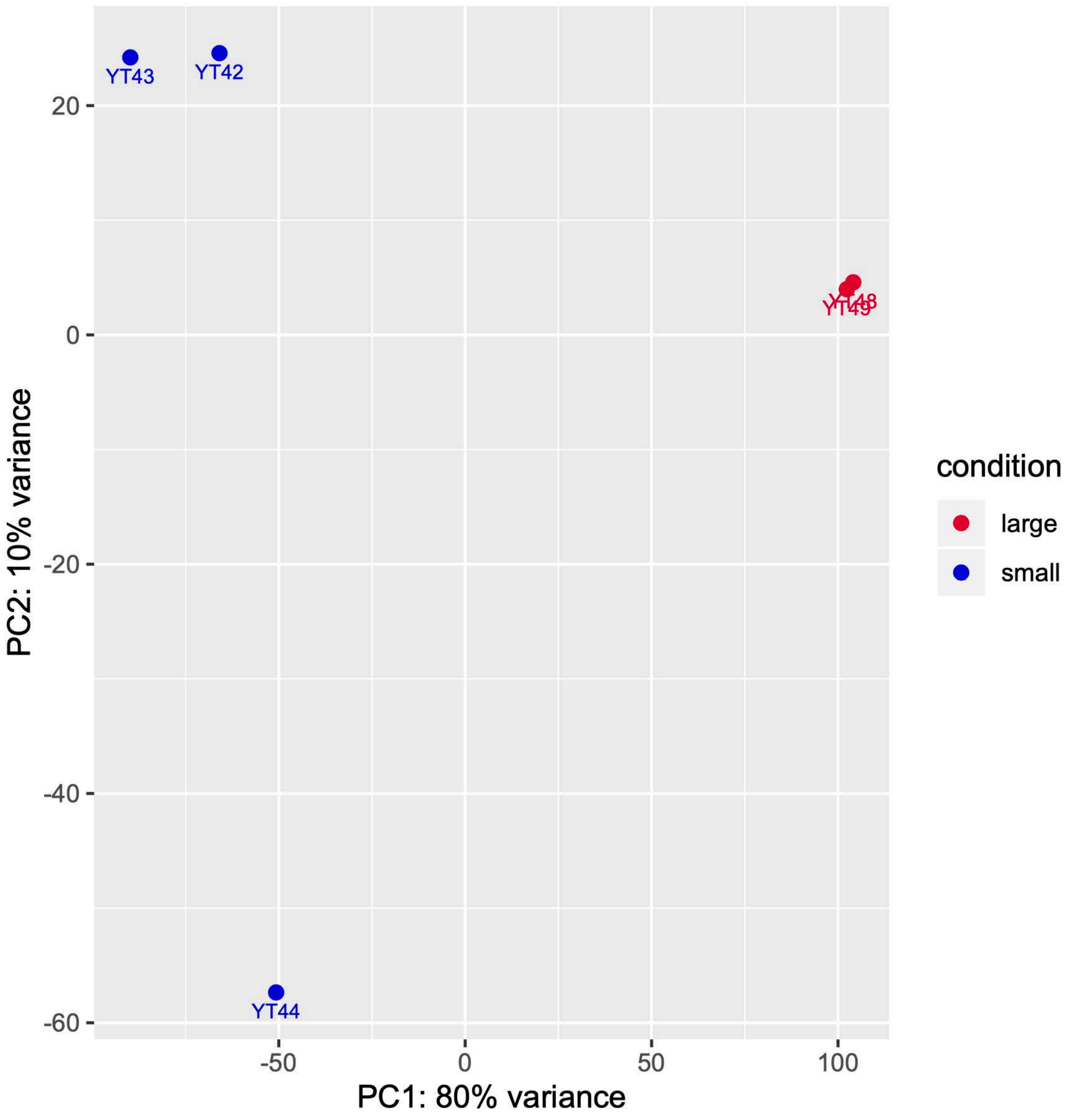


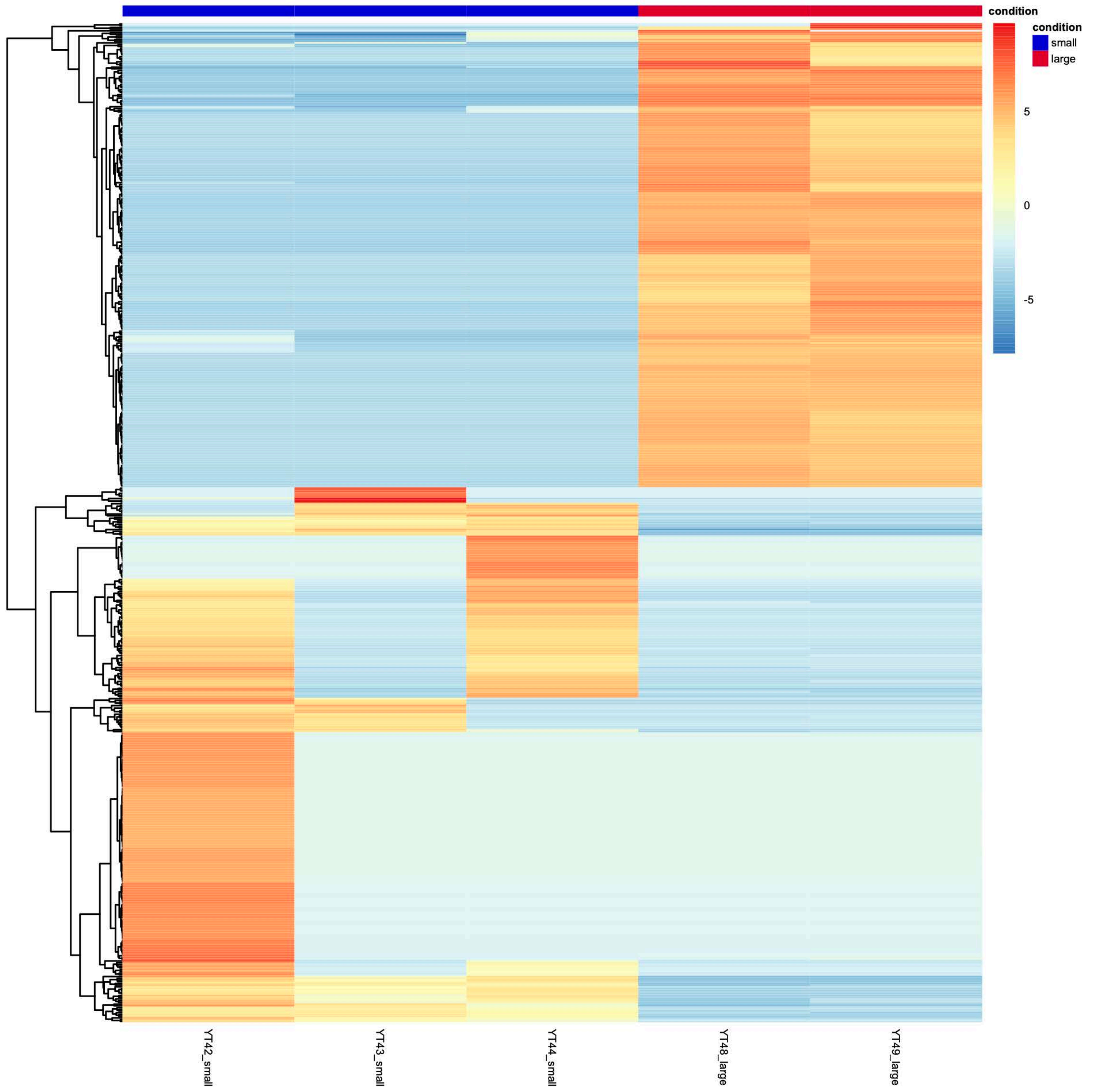

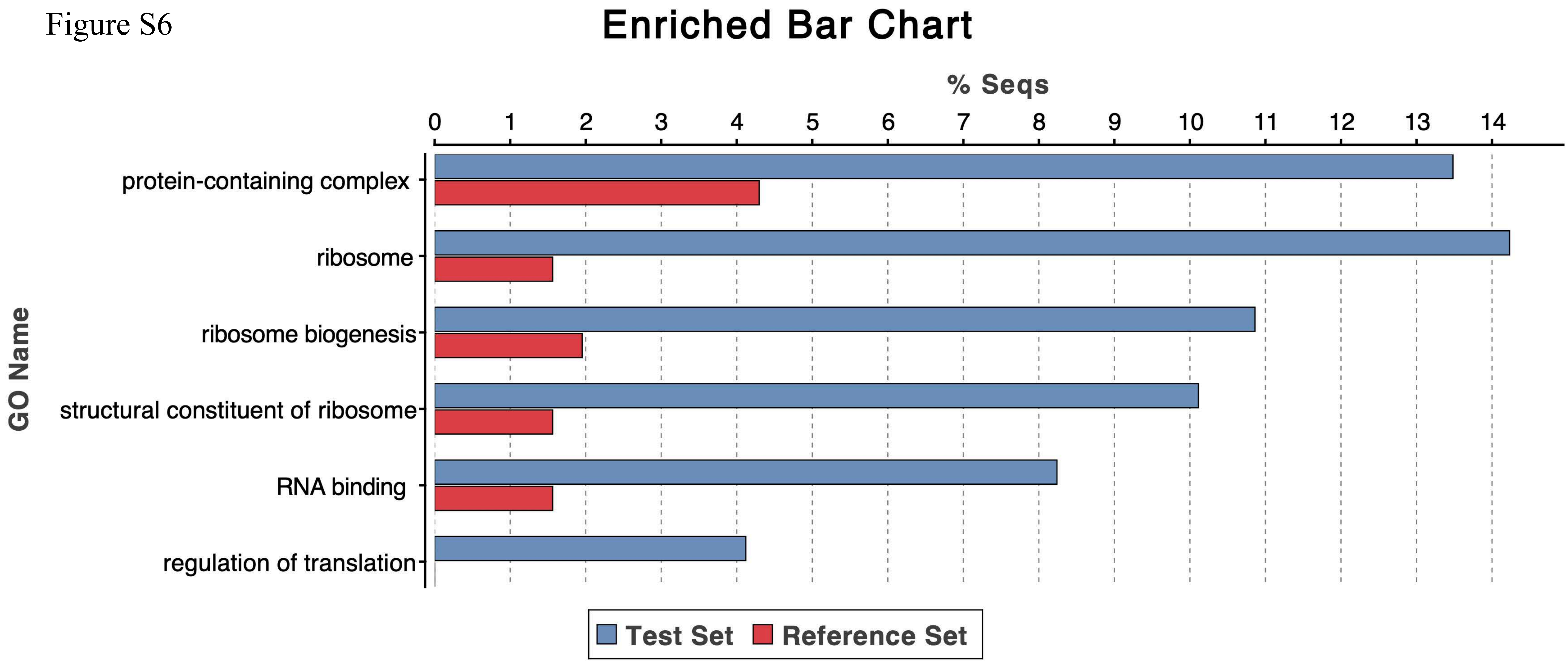

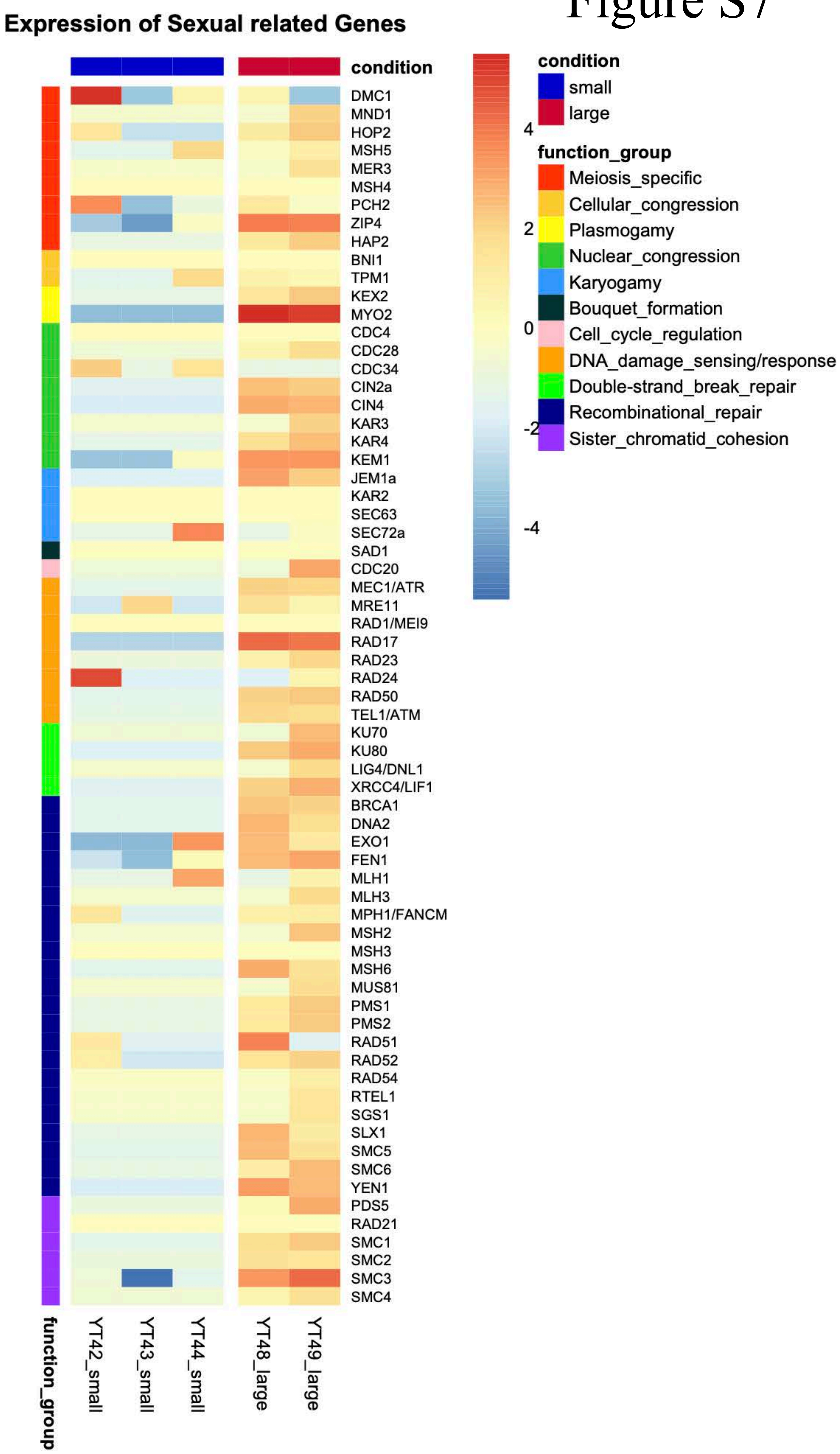

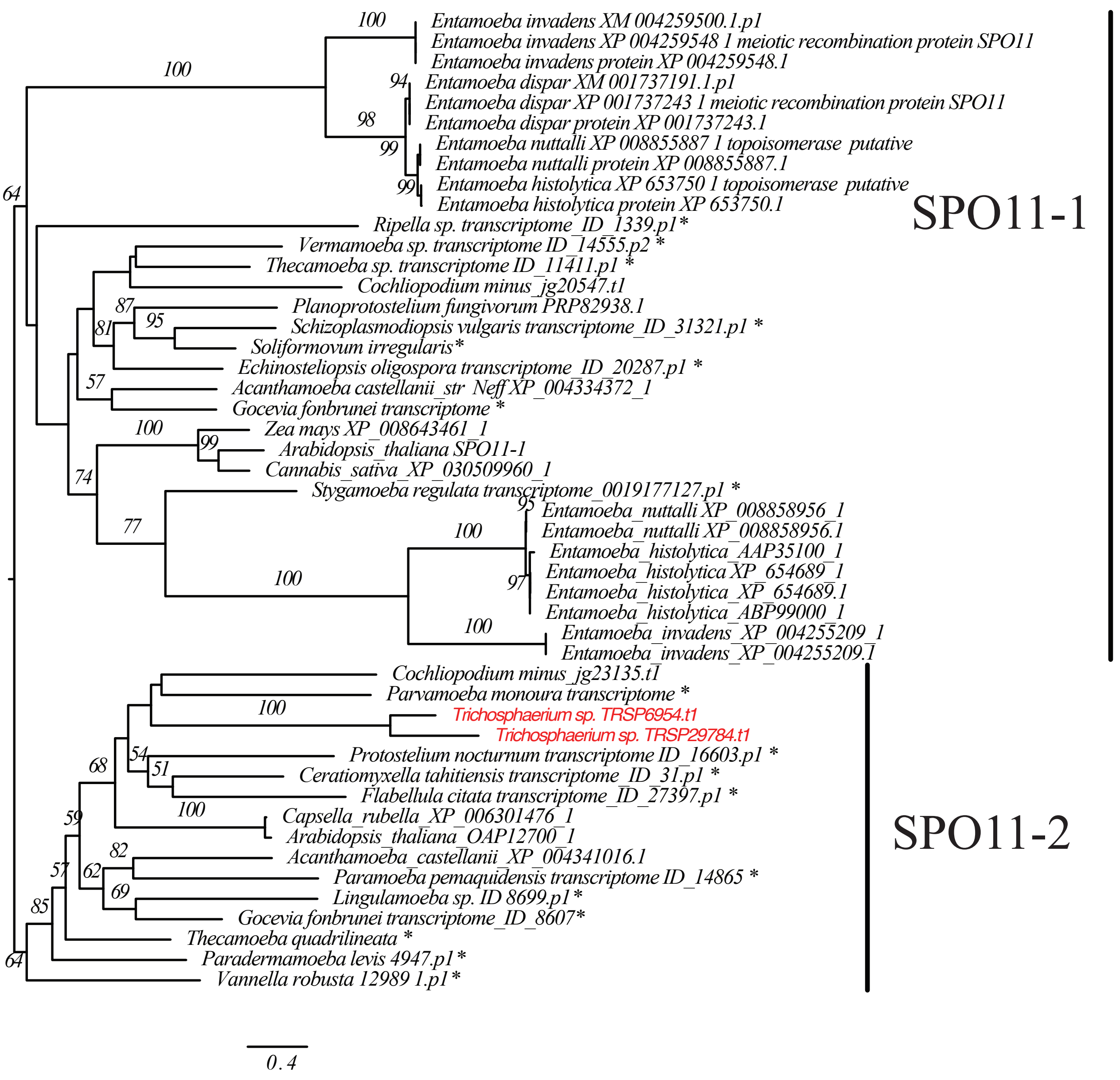
